# Genome-wide association studies unveil major genetic loci driving insecticide resistance in *Anopheles funestus* in four eco-geographical settings across Cameroon

**DOI:** 10.1101/2024.06.17.599266

**Authors:** Mahamat Gadji, Jonas A Kengne-Ouafo, Magellan Tchouakui, Murielle J Wondji, Leon Mugenzi, Jack Hearn, Boyomo Onana, Charles S. Wondji

**Author notes:** Corresponding authors **E-mails:** (MG) and (CSW).

## Abstract

Insecticide resistance is jeopardising malaria control efforts in Africa. Deciphering the evolutionary dynamics of mosquito populations country-wide is essential for designing effective and sustainable national and subnational tailored strategies to accelerate malaria elimination efforts.

Here, we employed genome-wide association studies through pooled template sequencing to compare four eco-geographically different populations of the major vector, *Anopheles funestus,* across a South North transect in Cameroon, aiming to identify genomic signatures of adaptive responses to insecticides. Our analysis revealed limited population structure within Northern and Central regions (*F_ST_*<0.02), suggesting extensive gene flow, while populations from the Littoral/Coastal region exhibited more distinct genetic patterns (*F_ST_*>0.049). Greater genetic differentiation was observed at known resistance-associated loci, resistance-to-pyrethroids 1 (rp1) (2R chromosome) and CYP9 (X chromosome), with varying signatures of positive selection across populations. Allelic variation between variants underscores the pervasive impact of selection pressures, with rp1 variants more prevalent in Central and Northern populations (*F_ST_*>0.3), and the CYP9 associated variants more pronounced in the Littoral/Coastal region (*F_ST_* =0.29). Evidence of selective sweeps was supported by negative Tajima’s D and reduced genetic diversity in all populations, particularly in Central (Elende) and Northern (Tibati) regions. Genomic variant analysis identified novel missense mutations and signatures of complex genomic alterations such as duplications, deletions, transposable element (TE) insertions, and chromosomal inversions, all associated with selective sweeps. A 4.3 kb TE insertion was fixed in all populations with Njombe Littoral/Coastal population, showing higher frequency of *CYP9K1* (G454A), a known resistance allele and TE upstream compared to elsewhere. Our study uncovered regional variations in insecticide resistance candidate variants, emphasizing the need for a streamlined DNA-based diagnostic assay for genomic surveillance across Africa. These findings will contribute to the development of tailored resistance management strategies crucial for addressing the dynamic challenges of malaria control in Cameroon.

**Author Summary:** Despite the widespread use of vector control tools to combat malaria in Cameroon, the disease burden remains high, particularly affecting children and pregnant women. This persistent burden is linked to intense resistance in malaria vectors, mainly driven by the overexpression of metabolic insecticide resistance genes. The evolutionary response of mosquito populations to both control interventions and agricultural environmental stimuli across Cameroon is not well understood. Understanding these dynamics is crucial for developing effective and sustainable strategies for malaria elimination country-wide.

Here, we performed a genome-wide survey of *Anopheles funestus* across four eco-geographic regions in Cameroon, revealing limited population structure between the northern and southern regions. For the first time in Cameroon, we observed the emergence and widespread of two known resistance-related loci, rp1 and CYP9 loci. Additionally, we identified both known and novel replacement polymorphisms, along with complex signatures of genomic alterations such as large insertions and duplications, linked to selective sweeps. Notably, a 4.3kb structural variant was completely fixed in all regions, while the *CYP9K1* resistant allele (A454A) was fixed only in the littoral/coastal region but remained under selection elsewhere highlighting the importance of designing a tailored resistance management strategies crucial for addressing the dynamic challenges of malaria control in Cameroon.

## Introduction

Malaria, caused by the *Plasmodium* parasite and transmitted by *Anopheles* mosquitoes, remains a major global health challenge with over 249 million cases and 608,000 deaths in 2022 [1]. In the African region alone, malaria contributed to 233 million cases and claimed the lives of 580,000 individuals in the same year, with children under 5 the most vulnerable to this disease [1].

Similar to other endemic regions, the fight against malaria in Cameroon relies heavily on insecticide-based interventions, such as insecticides treated nets (ITNs) and indoor residual spraying (IRS), which target the major mosquito vectors such as *Anopheles gambiae* and *Anopheles funestus* [2]. For the past years, the National Malaria Control Programme (NMCP) of Cameroon and their partners have invested much efforts to drive malaria towards elimination particularly through its National Strategy for the Health Sector, as well as the National Strategic Plan to Fight Against Malaria between 2019-2023 and 2023-2027 [3]. This was evidenced by the substantial increase in coverage of long-lasting insecticidal nets (LLINs) from 21% to 59% between 2011 and 2018 [3]. Additionally, the Cameroon government, under the "STOP MALARIA" initiative, has recently initiated its fourth LLIN distribution campaign. This campaign aims to distribute 16,756,200 LLINs in three phases, targeting a total of 27,740,035 Cameroonians. The distribution occurred in different regions, with specific types of nets provided based on regional characteristics, including standard LLINs, LLINs + PBO (an inhibitor of P450s enzymes associated with metabolic resistance), and new-generation dual-AI LLINs (Interceptor G2) [4,5]. Despite concerted efforts of the government and the decline of malaria incidence in Sub-Saharan African in the past decades [6], malaria still remains endemic in Cameroon, contributing to approximately 6.46 million cases and 9,000 deaths in 2022 [1].

Unfortunately, the major malaria vector *Anopheles* species have evolved resistance to all four classes of insecticides (pyrethroids, carbamates, organochlorines and organophosphates), particularly pyrethroids used in bed net impregnation. This resistance has emerged due to the extensive and widespread deployment of vector control tools, combined with the use of pesticides in the agricultural environment [7]. Consequently, insecticide resistance in malaria vectors poses a major threat to the effectiveness of current and future control strategies. The underlying mechanisms of insecticide resistance in these malaria vectors include target site knock-down resistance evidenced in *An. gambiae* sl and broadly absent in *An. funestus* [8,9], cuticular resistance [10], behavioural resistance [11] and metabolic resistance. Metabolic resistance acts through metabolism or sequestration of insecticides by metabolic enzymes majorly the cytochrome P450s, Glutathione S transferases and Carboxylesterases clusters [12–14]. However, there are substantial variations in the resistance patterns and its underlying mechanisms across different African regions including Cameroon [15–17]. Such resistance was mainly linked to the over-activity of metabolic resistance genes [15–17].

Population genomics plays a crucial role in identifying the molecular mechanisms underlying resistance dynamic and in understanding and predicting their spread in different populations. A population genomic approach is particularly useful when considering the population structure of the species throughout its range. Transitions between biomes and environmental heterogeneity in *Anopheles* species are frequently linked to local adaptation and ecological divergence [18–20]. Additionally, selection pressures related to pollutants, particularly those specific to certain environments, may confer environments specific signals of local adaptation linked to ecology [21,22]. For example, diverse agricultural practices have been correlated with variations in the frequencies of resistance mechanisms in *An. gambiae* s.l in Côte d’Ivoire [18] and in Cameroon [23].

Temporal Genome-Wide Association Study (GWAS) across Africa revealed notable genomic changes in *Anopheles funestus*, including the emergence of P450s-based loci (QTL rp1 and CYP9) with selective sweeps and reduced genetic diversity [15,17]. Mosquito populations are often considered to be homogenous across a country leading control programs to roll out the same control intervention nation-wide [2,24]. However, past evidence suggests that this one size fits all may not reflect the complexity of the genetic structure at the scale of a single country such as Cameroon where there are strong eco-climatic contrasts from south to north [25,26]. Indeed, in Malawi, it was shown that there was a significant variation of gene expression in relation to insecticide resistance in *An. funestus* from the southern part to the north [27]. The extent of such variation remains unknown and the detection of genomic signatures of such countrywide variation remain limited. At a time when most NMCPs are opting to use a sub-national tailoring approach to better deploy interventions to ensure that control tools can best match the profile of the local populations [28], it is essential to define nationwide patterns of gene flow, signature of major genomic drivers of resistance to better inform decision on which tool to deploy in different regions.

Consequently, investigating the spatial dynamics of adaptive response and the role of selection pressures among natural *Anopheles funestus* populations across Cameroon is crucial for the successful development and deployment of alternative control intervention tools that will help advance malaria elimination in the country by 2030. To fill this gap in knowledge, we conducted a comprehensive country wide pooled template (PoolSeq) genome wide association study (GWAS) of *An. funestus* mosquitoes sampled from four eco-geographical sites across Cameroon, a high burden, high impact malaria country [1].

Subsequently, we examined population structure and explored genomic differentiation across the whole genome of *An. funestus* in order to pinpoint distinctive signatures indicative of evolutionary selection spanning the potential insecticide resistance or novel loci. Furthermore, we conducted a thorough examination of replacement polymorphisms and signatures of complex genomic alterations associated with the selective sweeps to enhance our understanding of the evolutionary dynamics in these regions.

## Results

### Overview of alignment and coverage metrics

Alignment of PoolSeq GWAS produced 150×2bp reads ranging from 166 million for Gounougou population to 240 million reads for Elende population with a mean of 197 million reads. Filtering them according to read pairing, sequence quality, and mapping quality allowed successful mapping of reads on the *An. funestus* FUMOZ genome with mapping rates ranging from 29% for Njombe population to 90% for Gounougou population with a mean mapping rate of 65.75%. Most of the mapped reads aligned in pairs (>98%) and less than 8% of the reads for each sample were singletons. The mean coverage was homogeneous across all the samples, except for the sample collected in Njombe (∼25x < to the targeted coverage). This low mapping rate can negatively impact the downstream analysis, as certain genomic regions are underrepresented, skewing analyses like comparative genomics between this Njombe population and others. The mean coverages ranged from 25.70x for the Njombe population to 90.21x for the Tibati population, with a total mean coverage of 62.33x (S1 and S2 Table).

### Population structure

Pairwise *F_ST_* values were calculated among *An*. *funestus* populations collected in the four eco-geographical settings and presented in a correlation heatmap (Fig 1a). Lower *F_ST_* values were observed between *An. funestus* from Gounougou and Tibati (0.003) but stronger between Njombe and Elende (0.01), indicating extensive gene flow within the former and limited gene flow within the latter populations. Comparisons between Gounougou versus Njombe and Elende yielded slightly higher *F_ST_* estimates, ranging from 0.04 to 0.05. Tibati versus Njombe and Elende showed *F_ST_* values ranging from 0.02 to 0.04, suggesting limited gene flow between populations from Northern to Central and Littoral/Coastal regions of Cameroon. High *F_ST_* values were recorded between all four Cameroon populations and the reference strains FANG and FUMOZ (ranging from 0.21 to 0.35), supporting little or restricted gene flow between Central and Southern African *An. funestus* populations as previously reported [15] although this difference could also be due to inbreeding in the lab strains.

**Figure 1:**
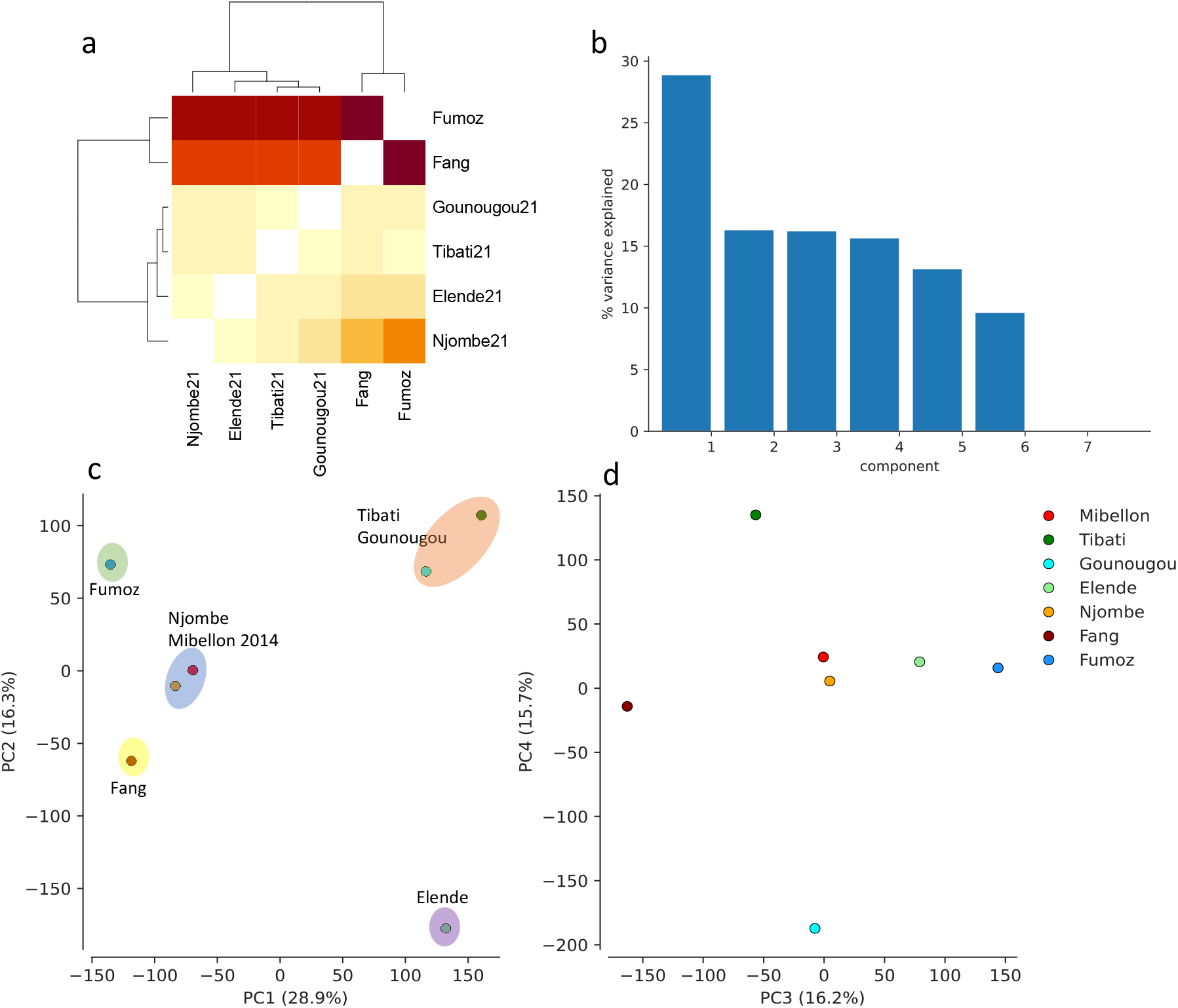
Genetic differentiation and population structure among *An. funestus* across Cameroon using pairwise *F_ST_* and PCA approaches. a: Pairwise *F_ST_* correlation heatmap between *An. funestus* collected in four different eco-geographical settings across Cameroon; b: PCA components capturing the major signals of *An. funestus* population subdivision across Cameroon; c and d: PCA showing the genetic structure of *An. funestus* populations across Cameroon with Fig 1c showing that the majority of the variance observed was capture by PC1 1 and PC2 compared to PC3 and PC4 in Fig 1d.

To further confirm the minimal observed population structure, historical relationships and genetic structure among populations were analyzed using Principal Component Analysis (PCA). PCA clearly separated Northern populations (Gounougou and Tibati) from Central and Littoral/Coastal populations (Njombe and Elende), while the clustering of FANG and FUMOZ populations from Southern Africa corresponds with their geographical location, albeit showing genetic divergence (Fig 1c). PCA supports the previous findings by revealing a clear divergence between *An. funestus* populations from Northern regions and those from Central and Littoral/Coastal regions (Fig 1c), with the majority of variance supported by PC1 (Fig 1b). It clearly separated populations into three main clusters (Fig 1c). Northern populations (Gounougou and Tibati) were genetically more related to each other, while the Elende Central population formed a distinct cluster, separate from both Northern and Littoral/Coastal populations. Notably, the Mibellon 2014 population, despite its Northern location near Tibati (<100 km), clustered with the Njombe Littoral/Coastal population from a different region, suggesting potential genetic similarities between these populations. Mibellon is located more to the west before the Adamawa mountainous chain. This positioning could explain its differences from Tibati and its similarities with Njombe. These observations imply the presence of region-specific advantageous variants associated with the evolution of insecticide resistance, warranting further detailed investigation.

### Windowed measures of pairwise *F_ST_* population genetic differentiation

Windowed Interpopulation *F_ST_* comparisons among our four natural *An*. *funestus* populations were conducted to quantify the degree of genetic variation between them. Globally, high *F_ST_* values (>0.2) were observed across chromosomes for all relevant pairwise comparisons, including populations from the Northern to Central and Coastal regions of Cameroon. Notably, these signals were prominent around the known quantitative trait locus (QTL) rp1 (chromosome 2) and CYP9 (Chromosome X) resistance-associated loci (Fig 2).

**Figure 2:**
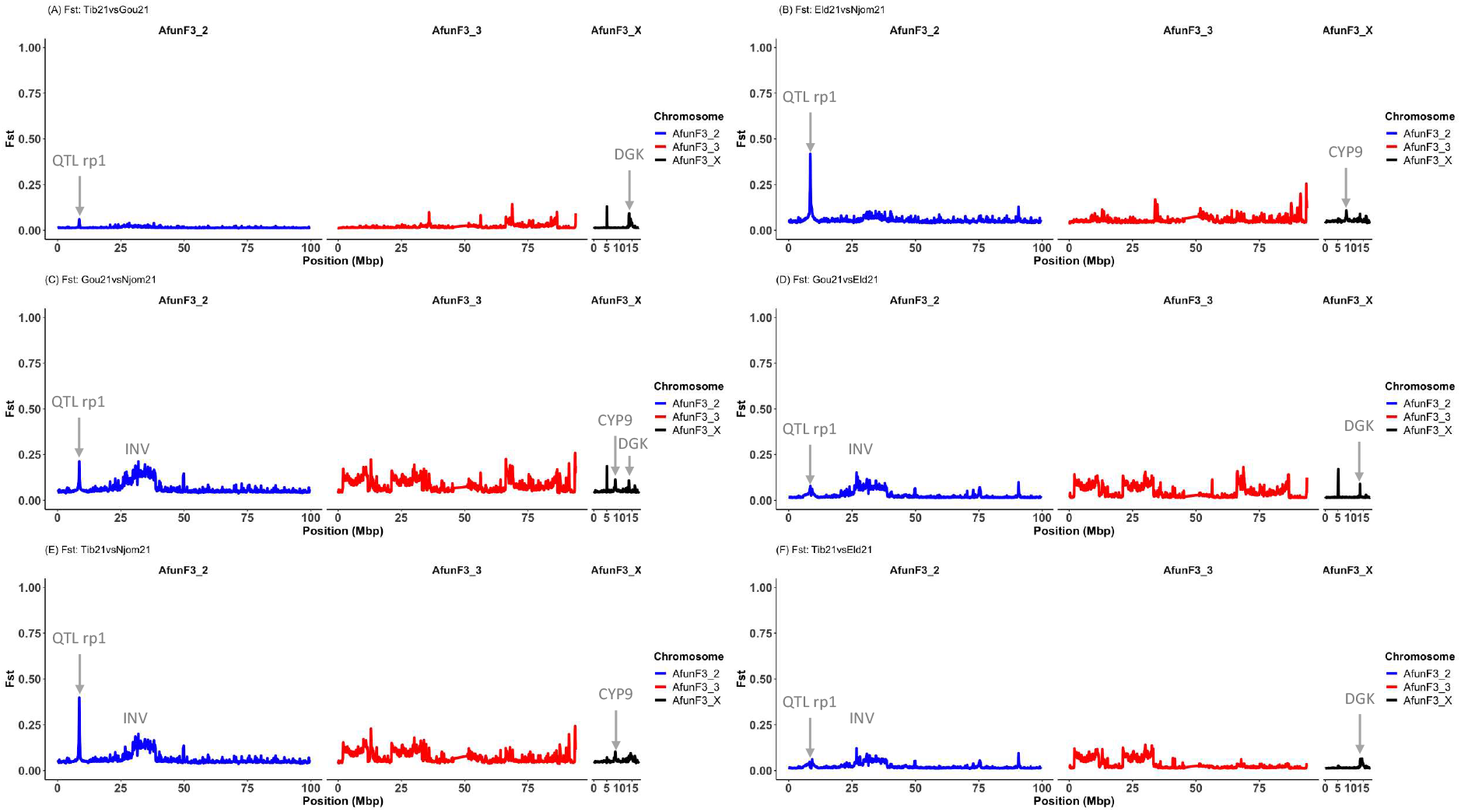
Pairwise *F_ST_* genetic differentiation among *An*. *funestus* populations in four eco-geographical settings across Cameroon. rp1 QTL, standing for Quantitative Trait Locus resistant to pyrethroid 1, denotes the genomic region responsible for 87% of observed pyrethroid resistance in some *An. funestus* [30]. Additionally, DGK refers to Diacylglycerol kinase (*AFUN020012*), while CYP9 designates the cytochrome 9 cluster housing the metabolic resistance gene *CYP9K1*. INV indicates region affected by chromosomal inversion.

Key findings include the presence of low genetic differentiation (*F_ST_* ∼ 0.06) between Gounougou and Tibati populations at the CYP6 locus only (Fig 2A). Comparisons between Elende and Njombe populations revealed emergence of both loci with a stronger genetic differentiation around the rp1 locus (*F_ST_* = 0.47), but lower at the CYP9 (*F_ST_* = 0.1) locus suggesting potential differences in insecticide resistance mechanisms (Fig 2B). Similarly, between Gounougou and Njombe populations, a peak of divergence emerged at both rp1 (*F_ST_* = 0.23) and CYP9 (*F_ST_* = 0.09) loci still stronger at the rp1 locus (Fig 2C). Gounougou versus Elende comparison only evidenced the emergence of rp1 locus with lower *F_ST_* value (*F_ST_* <0.1) compared to other comparisons (Fig 2D). Comparison between Tibati and Njombe *An. funestus* populations showed similar pattern as for Elende and Njombe comparison with a high signal of differentiation at both the rp1 (*F_ST_* = 0.44) and the CYP9 (*F_ST_* = 0.09) loci (Fig 2E). When comparing Tibati versus Elende populations, a weak sign of genetic differentiation is seen only at the known rp1 resistance-associated locus (*F_ST_* = 0.02) (Fig 2F). All comparisons except Tibati versus Gounougou, indicated the emergence of an extensive regions of around 6 Mbp, with high *F_ST_* values but no fixed differences on autosome 2 (Fig 2B-F). This region likely exhibits a pattern characteristic of complex chromosomal rearrangement, potentially involving chromosomal inversion. Exploration of this region revealed several overlapping inversions of significant length, indicating a more complex genetic picture than previously understood using microsatellite markers [29]. Moreover, all comparisons highlighted divergence on the X chromosome around 13.8 Mbp at varying *F_ST_* values, coinciding with diacylglycerol kinase (DGK) locus (*AFUN020012*). This suggests that while this locus has not been directly linked to insecticide resistance in malaria vectors previously, it could be under selection pressure. The comparison of all these populations to the fully multiple insecticide-susceptible FANG reference strain revealed a stronger selection pressure acting on both the rp1 and CYP9 regions with varying *F_ST_* values: For the rp1 locus, in Tibati (*F_ST_* = 0.36), Gounougou (*F_ST_* = 0.23), Elende (*F_ST_* = 0.38) and Njombe (*F_ST_* = 0.29) and for the CYP9 locus, in Tibati (*F_ST_* = 0.19), Gounougou (*F_ST_* = 0.19), Elende (*F_ST_* = 0.19) and Njombe (*F_ST_* = 0.29) (S1 Fig). Knowing that there was no sign of genetic variation associated to insecticide resistance in Mibellon in 2014, additional genomic comparisons using it as a control confirmed major genomic changes in these *An. funestus* populations across Cameroon. The findings obtained support the previous observations with consistent patterns noted around the differentiated regions. Indeed, pairwise comparisons between Mibellon 2014 and all *An. funestus* populations revealed that the rp1 locus is stronger in Elende and Tibati compared to Gounougou and Njombe, with *F_ST_* values ranging between 0.19 for Gounougou and 0.38 for Elende population while the CYP9 peak was found at higher *F_ST_* (0.29) in Njombe population (Fig 3) indicating that the CYP9-based mechanism is stronger in Littoral/Coastal *An. funestus* populations compared to others. These findings validate previous observations and highlight regional variations in insecticide resistance mechanisms across Cameroon.

**Figure 3:**
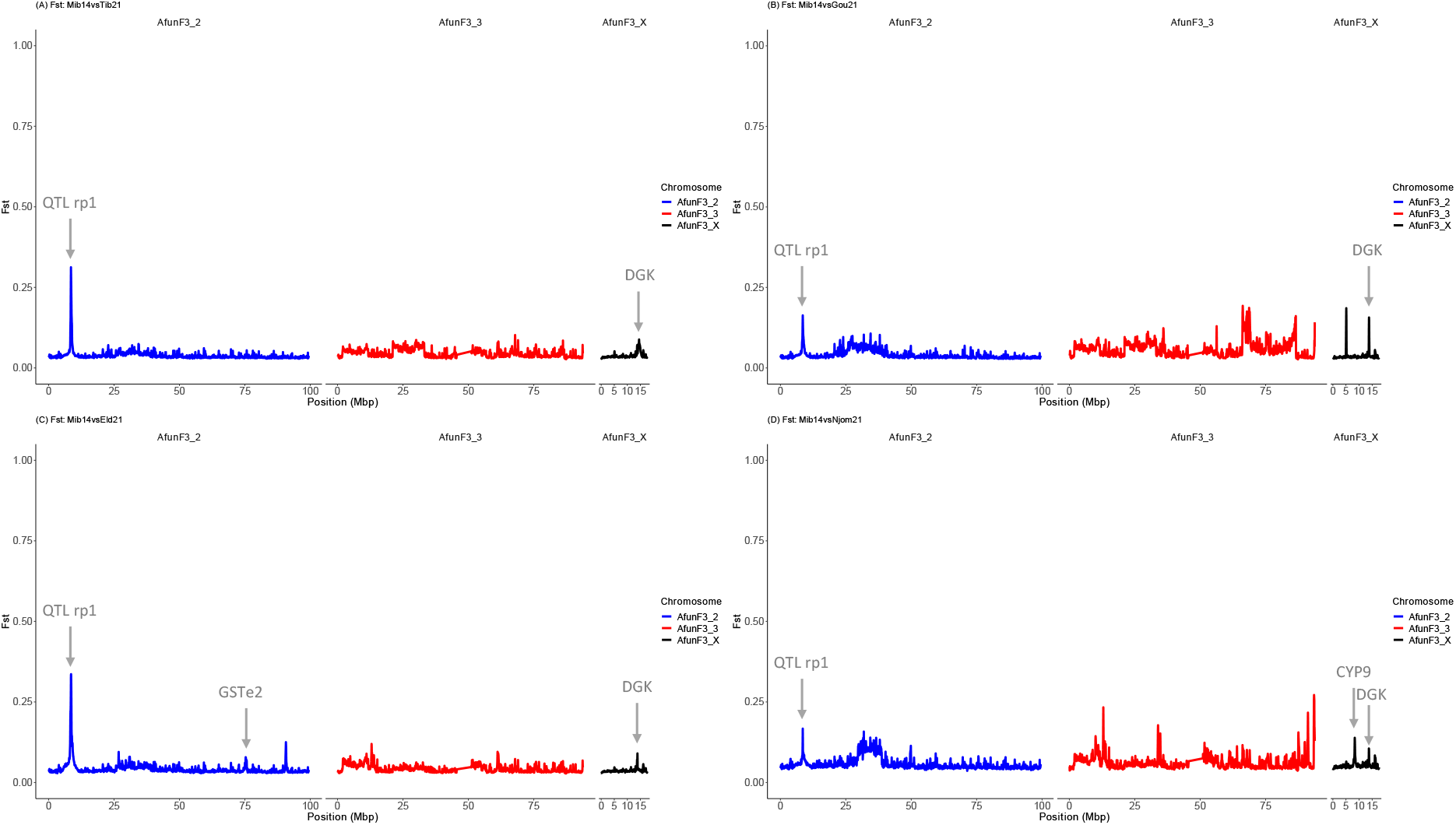
Pairwise *F_ST_* genetic differentiation between *An. funestus* population from four eco-geographical settings and Mibellon 2014 population.

### Population genomics

Analysis of genetic differentiation among *An. funestus* populations from the four eco-geographical localities allows the identification of two known resistance-associated genomic regions spanning the rp1-QTL and the CYP9 loci. To find out whether these genomic regions are under positive selection, we conducted Tajima’s D and theta π genetic diversity analyses spanning both loci. Tajima’s D distributions were calculated for windows of 50kb assigned to each chromosome arm (AfunF3_2, AfunF3_3, and X) for each population. Median Tajima’s D values were consistently negative across all chromosomes in all populations suggesting population expansion, or demographic events acting on these chromosomes. Lower Tajima’s D values were observed in Tibati, Gounougou, and Elende populations (D < -2) compared to Njombe population (D > -1) but were higher (D>0) in southern African laboratory strains (Fang and Fumoz) than in other populations (Fig 4). Interestingly, lower median Tajima’s D values were observed across the sex-linked X chromosome in all the four populations, with the lowest median values in Tibati (D = -2.15), followed by Gounougou (D = -2.05), Elende (D = -1.98), and Njombe (D = -0.9) populations. This observation potentially indicates differences in selective pressures or demographic histories among these populations acting on X chromosome.

**Figure 4:**
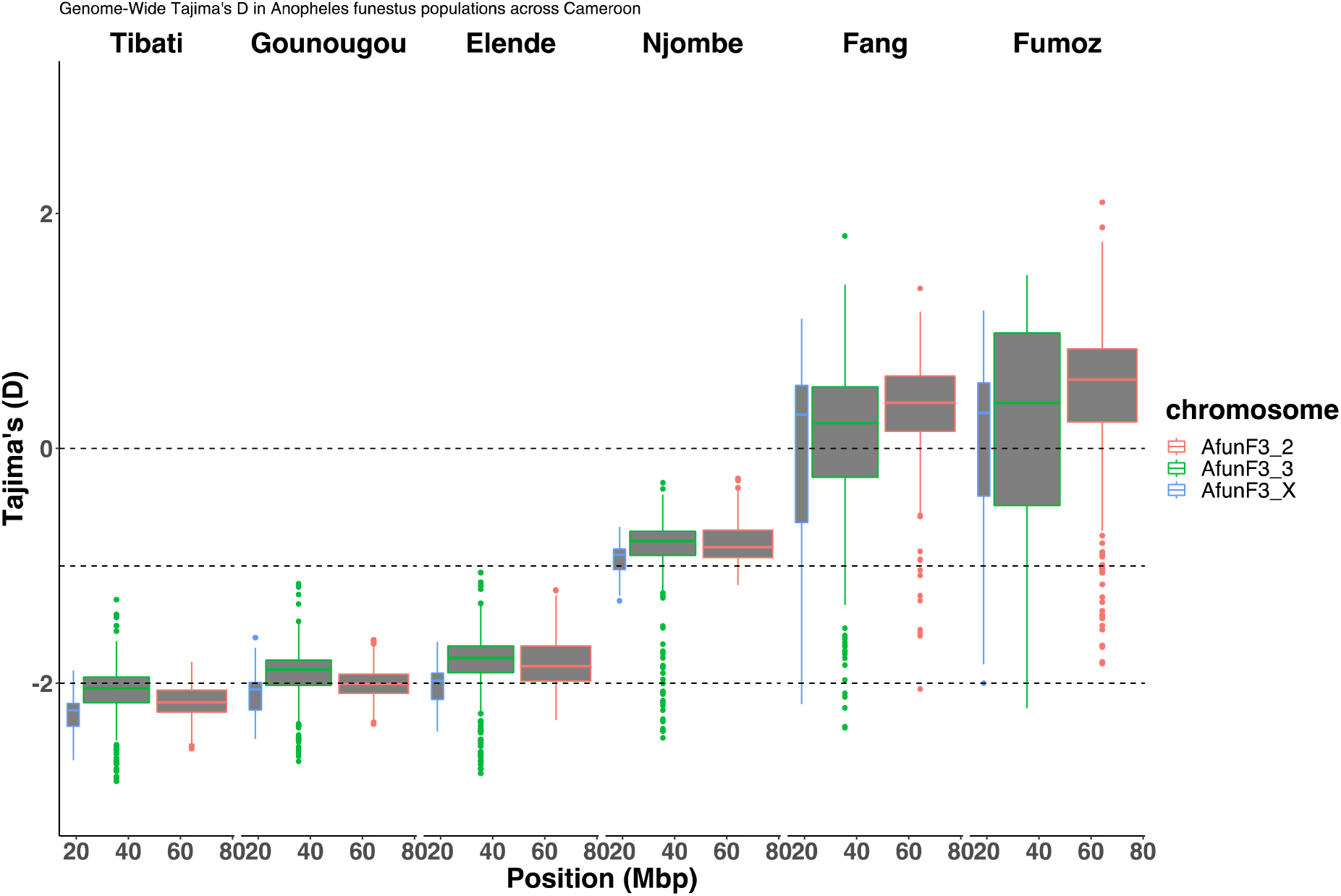
Genome-wide Tajima’s D calculated in overlapping windows of 50kb in *An. funestus* across Cameroon.

Zooming in around both resistance-associated loci (CYP6 and CYP9) for a finer resolution in 1kb overlapping windows revealed a valley of varying negative median Tajima’s D values spanning the entire rp1 locus, notably more pronounced in Elende and Tibati populations compared to the Gounougou and Njombe population (Fig 5A), which exhibited a higher median Tajima’s D. This suggests compelling evidence of a strong selective sweep across the entire rp1 QTL cluster (indicated by the red arrow) with a consistent valley observed in Elende and Tibati populations compared to the other populations aligning with their genetic differentiation patterns. FANG and FUMOZ populations appear to be in equilibrium (Fig 5A). Furthermore, the selective sweep at the rp1 locus was supported by a pattern of reduced genetic diversity mainly observed in Elende and Tibati *An. funestus* populations compared to Gounougou and Njombe populations which have similar pattern (Fig 5C). This demonstrates a consistent valley of selection strongly acting in Elende and Tibati populations than in Gounougou and Njombe in line with their genetic differentiation profile.

**Figure 5:**
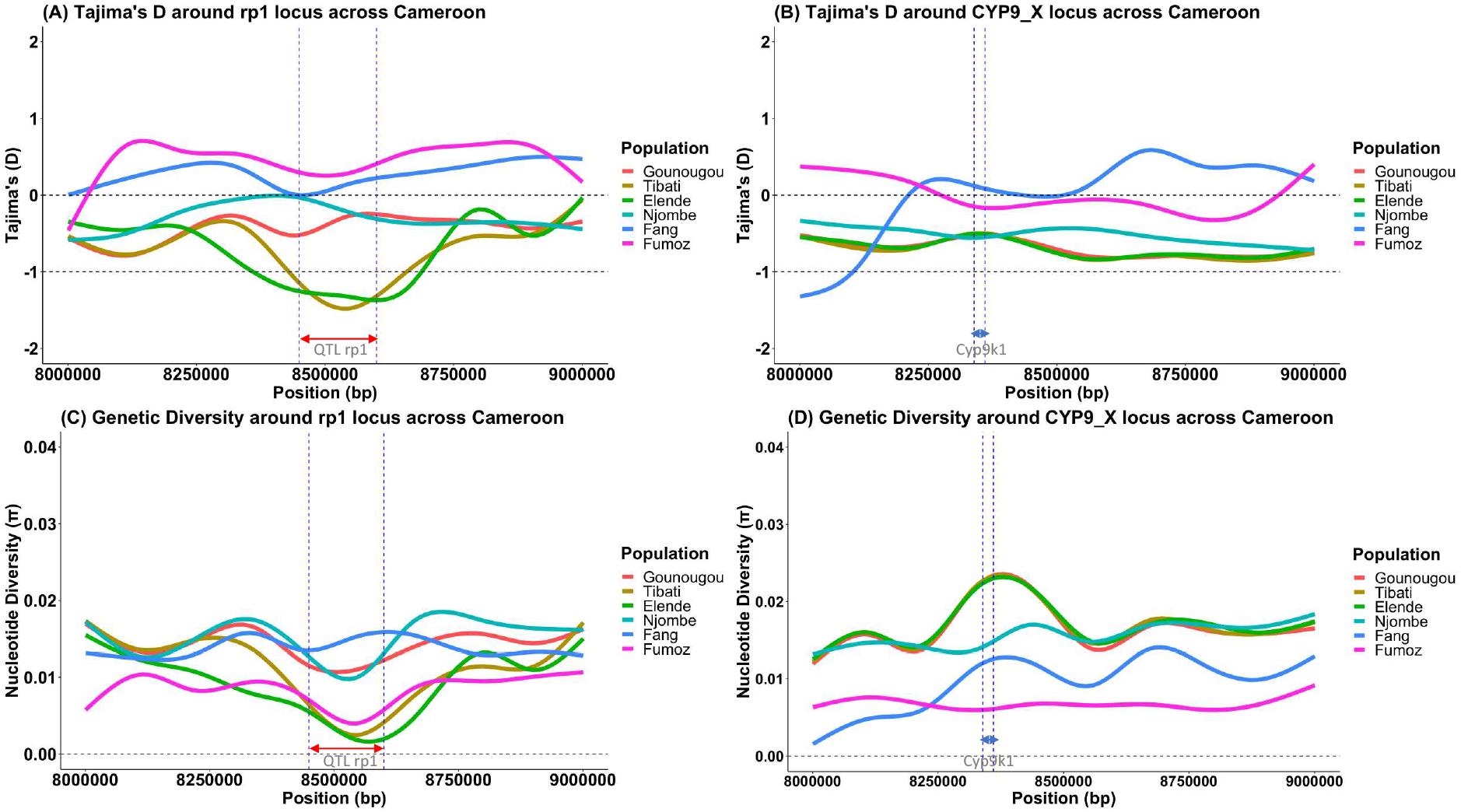
Selective sweeps around the rp1 QTL and CYP9 loci in *An. funestus* across Cameroon.

Concerning the CYP9 locus, a consistent pattern emerged, with Tajima’s D exhibiting negative and lower values specifically in the region spanning the *CYP9K1* gene (indicated by the red arrow in Fig 5B). Notably, Elende, Tibati, and Gounougou populations displayed lower Tajima’s D values (D ∼ -1.8) compared to Njombe (D ∼ -1) and laboratory strains FUMOZ and FANG, which were closer to equilibrium. This observation suggests a widespread and intensive selective sweep within these three populations, particularly targeting the CYP9 cluster and specifically the *CYP9K1* gene, a putative gene associated with insecticide resistance. Despite the overall low genetic diversity observed in all populations, a surprising reduction in diversity was noted in FANG and FUMOZ, probably due to inbreeding (Fig 5D) compared to the natural populations. It was also interesting to note that genetic diversity was more reduced in the Njombe population compared to others, suggesting that genetic variants driving the selection of the *CYP9K1* gene could be more impacted in Njombe than in other populations (Fig 5D). This nuanced and dynamic evolutionary history implies complex adaptive processes, warranting further investigation into the genetic mechanisms underlying local adaptation and/or insecticide resistance in these *An. funestus* populations around this sex-link X chromosome.

Additional analyses of minor and major allele frequencies were conducted at both loci across these populations, shedding light on the genetic diversity patterns. Notably, consistently low minor allele frequencies (<0.05) were observed flanking both the rp1 (S8A Fig) and CYP9 (S8B Fig) clusters in all populations, suggesting the presence of rare genetic variants associated with these loci and supporting the previously observed pattern of selective sweeps.

### Variants associated to insecticide resistance evolution in *An. funestus* across Cameroon

#### Candidate SNPs associated with insecticide resistance

A comprehensive scan for replacement polymorphisms led to the identification of 185,623 missense variants distributed across chromosomes. Filtering based on allele frequencies, coverage depth, p-values, and variants located around active sites or substrate binding pockets of relevant genes resulted in the selection of 45 non-synonymous single nucleotide polymorphisms (ns-SNPs) from multiple genes. Special emphasis was placed on missense variants spanning resistance-associated genomic loci, including putative resistance-associated SNPs or loci such as the rp2, rp3, Ace-1, GABA-based rdl mutations, GSTs cluster, and screening of the voltage-gated sodium channel (VGSC), extending beyond the loci detected in this study. Distinct patterns of SNP allelic frequencies were observed among populations, with Elende (Central), Tibati and Gounougou (Northern) exhibiting a similar pattern, contrasting with the observed pattern in the Njombe (Littoral/Coastal) population (Fig 6 and S3 Table). Notably, the most significant novel ns-SNPs are spread across the rp1 locus, predominantly located around the substrate binding pockets of the corresponding genes, such as *CYP6P2* (R346C), *CYP6P9a* (E91D, Y168H), *CYP6P5* (V14L), *CYP6AA2* (V3A, S498L, P35S), *CYP6P9b* (V392F, V359I), *CYP6AD1* (H35Y), *CYP6P4a* (H169R), *CYP6P1* (L13F) and Carboxylesterase *AFUN015793* (S307Y), were either highly prevalent or fixed in Elende, Tibati and Gounougou populations. In contrast, they were either absent or present at low frequencies in Njombe, FANG, and FUMOZ laboratory strains, with the exception of *CYP6P9a* (E91D), which was fixed in Njombe. Conversely, the most significant SNPs in the Njombe population were dispersed across different genes, featuring just one P450s-based ns-SNP, *CYP9K1* (G454A). This SNP was nearly fixed in the Njombe population, while Elende, Tibati and Gounougou exhibited them at lower frequencies (<50%).

**Figure 6:**
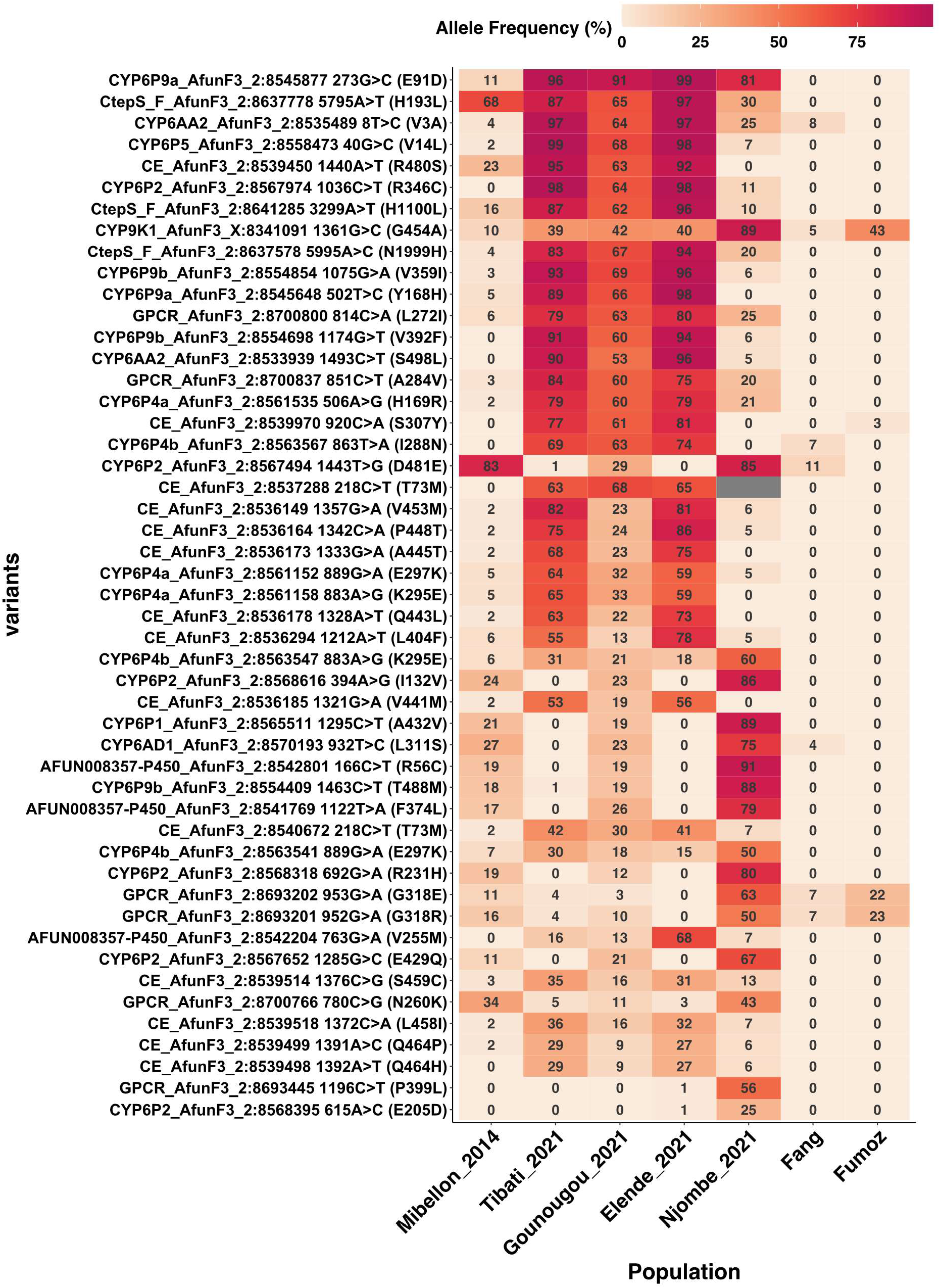
Heatmap showing the best replacement polymorphisms associated to insecticide resistance in *An. funestus* populations across Cameroon. Only the best non synonymous variants identified from frequency, depth, genotypes and p-value based-filtering.

These findings suggest that the single nucleotide polymorphisms (SNPs) surrounding the rp1 locus primarily correlate with insecticide resistance in Elende, Tibati, and Gounougou, in contrast to the Njombe population. In Njombe, the CYP9 (*CYP9K1*) locus appears to be more significant, exhibiting a fixed frequency (89.5%), while in other populations, it is present at moderate frequencies or is still under selection, with frequencies below 50% (Fig 6 and S3 Table). Additional SNPs identified at moderate to high frequencies, which flank the rp1 locus and are more prevalent in Njombe but present at lower frequencies in other populations, include novel variants. Among the novel SNPs are those on *CYP6P4b* (K295E, E297K), *CYP6P4a* (E297K, K295E), and *CYP6P4b* (I288N). Additional novel SNPs have been discovered on *CYP6P1* (A432V), cytochrome P450 *AFUN008357* (F374L, R56C), and CYP6AD1 (L311S). A SNP on a transcription factor gene, *AFUN019663* (TFIID, T16N) was fixed in all populations (>90%) but at very low frequency in FANG (8.7%), fully susceptible strain. Examination of target site knockdown resistance in these populations revealed no evidence of relevant SNPs in the VGSC capable of driving resistance (Fig 6). Replacement polymorphisms within the G protein-coupled receptor (GPCR) were detected at moderate to high frequencies, with some nearing fixation in Elende, Gounougou, and Tibati populations (GPCR A284V and L272I). In contrast, the Njombe population exhibited low to moderate allele frequencies for different GPCR polymorphisms (P399L, N260K, G318E, and G318R) (Fig 6). Although no hits were found around the rp2 and rp3 loci, scanning for missense variants within these loci revealed no significant SNPs associated with insecticide resistance. The polymorphisms observed were at low frequencies, with the highest variant exhibiting an allele frequency of 27.40% (S3 Table).

Upon examination of the GSTs gene cluster, we identified SNPs at moderate to high allelic frequencies that are present in all four populations. For instance, prominent SNPs were observed in the *GSTD1* gene (G191D and Y209F) and the *GSTT1* gene (A224T), with allelic frequencies ranging from 60% in the Njombe population to 92.7% in the Tibati population (Fig 7). Additionally, the well-known resistance-associated SNP on the *GSTe2* gene (L119F), which confers resistance to DDT and permethrin, was found at varying allelic frequencies across locations, ranging from 41.9% in Tibati to 62.6% in the Elende populations (Fig 7). This SNP was closely followed by the F120L mutation on the same gene, which exhibited varying frequencies across locations, ranging from 2.4% to 33.7%. Another SNP on the *GSTe2* gene (K146T), previously identified in the *An. funestus* population in the Democratic Republic of the Congo (DRC), was detected at low frequencies in northern and central populations but was completely absent in the Littoral/coastal population.

**Figure 7:**
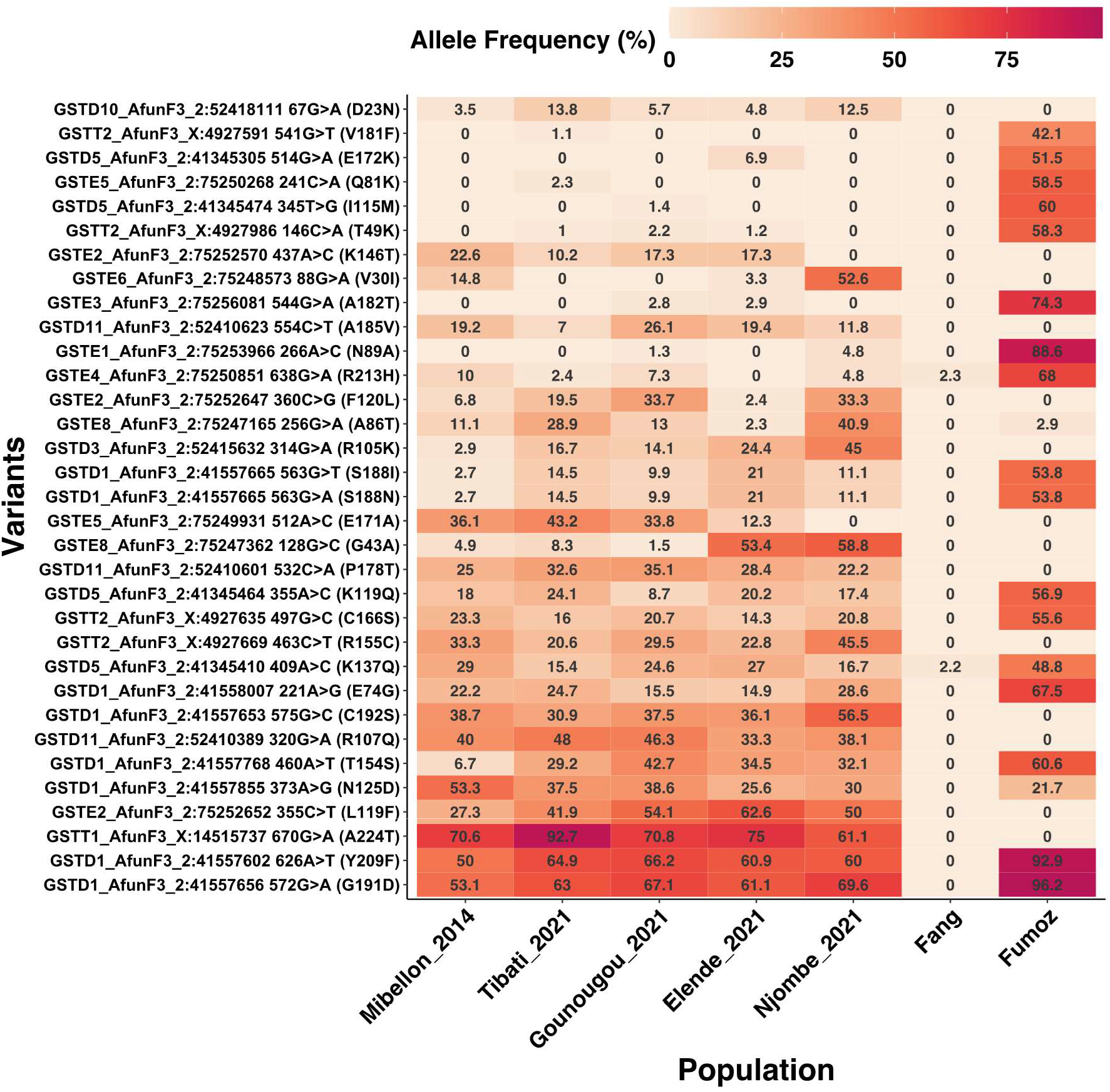
Non-synonymous polymorphisms in GST genes potentially associated with insecticide resistance in *An. funestus* across Cameroon.

Besides the two major CYP6 and CYP9-based selective sweeps, we examined target site insecticide resistance putative genes for evidence of relevant variants associated with insecticide resistance. Upon examination of the GABA receptor known to confer dieldrin resistance (rdl) in *An. funestus*, we identified three SNPs, with particular focus on two major ones. The first SNP, A296S, was observed at low frequencies in northern regions but was already fixed in central and littoral/coastal regions. The second SNP, T345S, was found to be fixed solely in the littoral region (Fig 8). Two SNPs were detected in the VGSC (D1156Y and L1988I) at very low frequencies (<11% and <17%, respectively) across all populations, with Njombe having the highest frequency for both SNPs (10% and 16.7%) while a single SNP was found on *Ace1* (AFUN011616), with a very low frequency in the Elende population (7.5%) and absence in other populations (Fig 8), suggesting that it has no role in insecticide resistance in *An. funestus* in all these localities across Cameroon.

**Figure 8:**
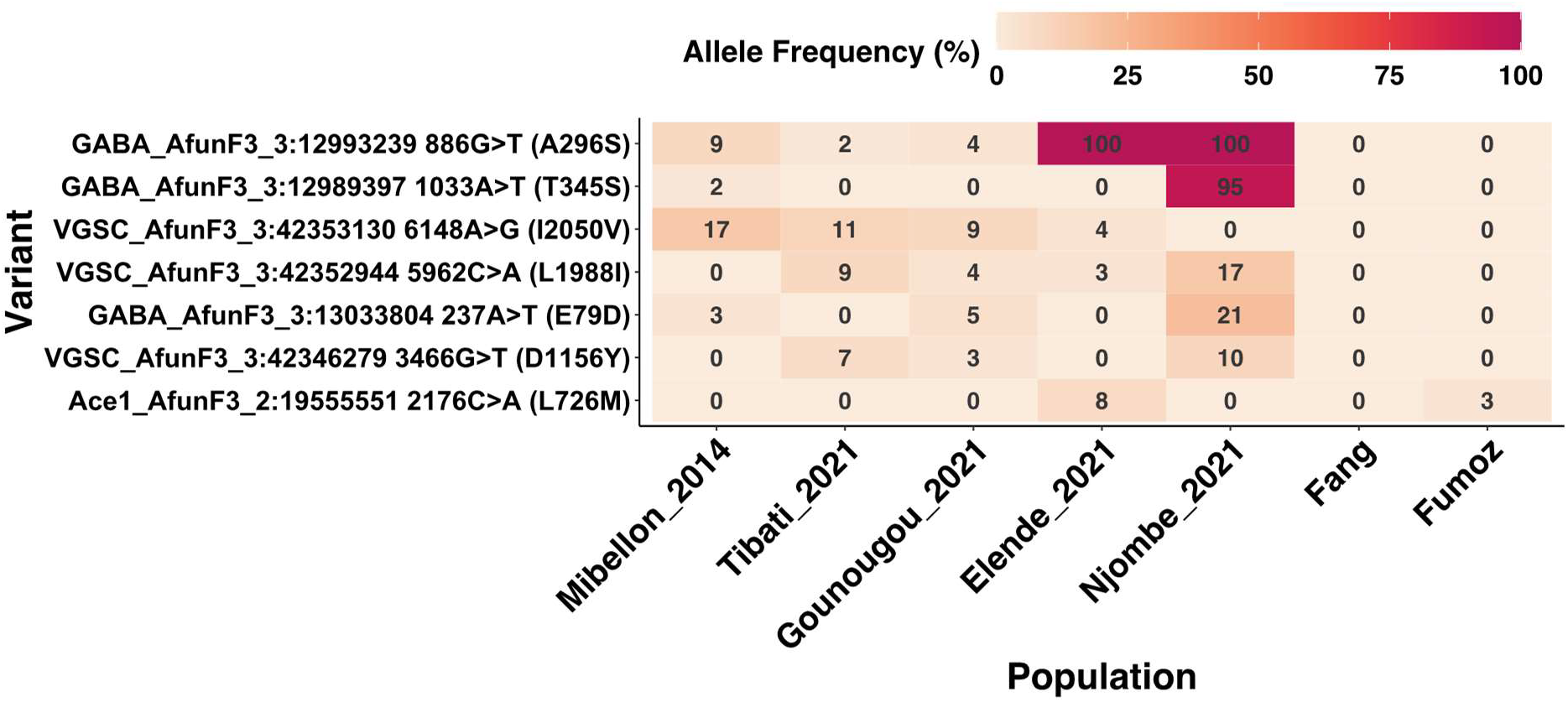
Non-synonymous polymorphisms within the target site genes potentially associated with insecticide resistance in *An. funestus* across Cameroon.

### Candidate signatures of complex genomic features associated to resistance evolution in *An. funestus* across Cameroon

A comprehensive large insertion calling was done across the rp1 and CYP9 loci using INSurVeyor tools and pertinent insertions were visualized in integrative genomic viewer (IGV). Our analysis revealed evidence of 26,724 insertions spread Genome-wide with around 233 spanning the entire rp1 and CYP9 loci. Of these, 2 large structural variants were pertinent and found in all the four populations but not in FANG and FUMOZ reference strains (S2, S3 Figs and S4 Table).

### Transposon insertions around the rp1 and CYP9 loci potentially associated to insecticide resistance in *An. funestus* across Cameroon

Mobile DNA sequences known as jumping genes play a crucial role in the dynamic nature of genomes, capable of relocating within the genetic material.

This study sheds light on a distinctive pattern involving the insertion of two large inserted transposable elements (TEs). One of these TE is found in the intergenic region between the CYP6P9b and CYP6P5 genes on chromosome 2R, specifically within the rp1 locus. The second one occurs upstream of the *CYP9K1* gene, in the vicinity of the CYP9 locus on the X chromosome. The initial transposable element (TE) is a 4.3 kb sequence previously detected in the Mibellon area of Cameroon. This sequence has been associated with pyrethroid resistance and reduced parasite infection in *An. funestus* populations across various regions of Cameroon [31]. The variant is inserted at position 8,556,409 and carries the insertion sequence “CCAAATGTACA.” Interestingly, this TE is fixed in all four studied localities (S2 Fig and S4 Table). The absence of this 4.3kb TE in the FANG susceptible reference strain further support its role in insecticide resistance. The robustness of this TE presence is supported by a high coverage depth and a consistent number of split reads, discordant reads, and supporting reads flanking both the left and right breakpoints. Additionally, the elevated scores (0.971/1 and 0.978/1 for the left and right breakpoints, respectively) surrounding these breakpoints further support the potential role of this TE in driving insecticide resistance in these populations (S4 Table).

On the other hand, in the upstream region of the *CYP9K1* gene, this study identified a second transposable element (TE) similar to the transposon detected between *CYP6P9b* and *CYP6P5*. This retrotransposon is inserted at position 8,338,432, characterized by an insertion sequence of “CAAATTTC” (S3 Fig and S4 Table). Notably, the pattern of this transposable element was more pronounced in the Njombe population compared to other populations, hinting at a contrasting distribution of this TE between the Northern, Central *An. funestus* populations, and the Littoral/Coastal populations of Cameroon (S3 Fig). Like the previously 4.3kb transposable element, this pattern was absent in the FANG susceptible reference strain. Unfortunately, the applied assembly approach encountered challenges in assembling the entire sequence, providing only around 830 bp with a pattern indicative of incomplete assembly represented by underscores ("_") (S4 Table). Despite the incomplete assembly, this transposable element was associated with consistent discordant and supporting reads, displaying a high coverage depth downstream that encompasses the *CYP9K1* gene. This observation suggests that this large transposable element could exert an impact on nearby genes, notably the *CYP9K1*, emphasizing the need for detailed investigations into its properties and potential implications in insecticide resistance.

### Candidate duplications, deletions and inversions around the rp1 and CYP9 loci associated to the selective sweeps in *An. funestus* across Cameroon

While large insertions were initially identified using INSurVeyor, we further investigated additional structural variants, including duplications, deletions, and inversions, utilizing smoove. Our focus was on structural variants with a length greater than 1kb, particularly those flanking the rp1 and CYP9 clusters, known to be under selection. Our analysis revealed a total of 324 structural variants distributed across both the rp1 and CYP9 genomic regions in *An. funestus* from the four natural populations, with variant counts of 94, 92, 86, and 52, respectively (S5 Table). These variants comprised tandem duplications, deletions, and complex chromosomal inversions. Notably, 28 of these variants emerged as particularly relevant, suggesting their potential contribution to *An. funestus* adaptive response to selective pressures induced by insecticide usage.

One noteworthy structural variant was a novel significant 2.4 kb duplication detected on chromosome 2 at position 8,560,624 and confirm by visualisation in IGV (Table 1, S5 Table and S4 Fig). This duplication spanned partial CYP6P4a and CYP6P4b paralogue genes known to be overexpressed in Ghana populations. Intriguingly, this duplication exhibited a heterozygote pattern in Elende, Tibati and Gounougou populations but was absent in Njombe population as well as FANG and FUMOZ laboratory strains, indicating ongoing selection of this duplication in the former populations (Table 1). Characterized by high coverage depth, consistent supporting reads, and sequence quality, this structural variant absence in the FANG reference strain suggested its potential role in insecticide resistance, warranting further investigation (Table 1, S5 Table). Another common duplication across all populations was a 3.5 kb variant partially spanning two carboxylesterases (*AFUN015793* and *AFUN015787*) within the rp1 locus. While fixed in Gounougou and Tibati, it presented as a heterozygote in Elende and Njombe populations, remaining absent in FANG (Table 1, S5 Table and S5 Fig).

**Table 1:** Overview of potential structural variations associated to insecticide resistance in *Anopheles funestus* population across Cameroon.

| Population | Chromosome | Start | End | SVs_type | SV_length | GT | Associated_Genes |
| --- | --- | --- | --- | --- | --- | --- | --- |
| Gounougou | AfunF3_2 | 8560624 | 8563109 | Duplication | 2.4kb | 0/1 | Spanning partial CYP6P4a and CYP6P4b |
| Gounougou | AfunF3_2 | 8527362 | 8534289 | Duplication | 6.9kb | 0/1 | Spanning entire CYP6AA1 and partial CYP6AA2 |
| Gounougou | AfunF3_2 | 8536226 | 8539803 | Duplication | 3.5kb | 1/1 | Spanning partial 2X Carboxylesterases |
| Gounougou | AfunF3_X | 8358864 | 8373842 | Duplication | 14kb | 1/1 | Spanning CYP9K1 and AFUN007545 intergenic |
| Gounougou | AfunF3_2 | 8560561 | 8562887 | Deletion | 2.3kb | 0/1 | Spanning partial CYP6P4a and CYP6P4b |
| Gounougou | AfunF3_2 | 8546959 | 8553509 | Deletion | 6.5kb | 1/1 | Spanning CYP9a and b intergenic region |
| Gounougou | AfunF3_X | 8347608 | 8377466 | Inversion | 29kb | 0/1 | Spanning CYP9K1 and AFUN007545 intergenic |
| Gounougou | AfunF3_X | 8356661 | 8405651 | Inversion | 48kb | 0/1 | Spanning entire AFUN007545 and next genes |
| Tibati | AfunF3_2 | 8560700 | 8563104 | Duplication | 2.4kb | 0/1 | Spanning partial CYP6P4a and CYP6P4b |
| Tibati | AfunF3_2 | 8536338 | 8539804 | Duplication | 3.5kb | 1/1 | Spanning partial 2X Carboxylesterases |
| Tibati | AfunF3_2 | 8560552 | 8562882 | Deletion | 2.3kb | 0/1 | Spanning partial CYP6P4a and CYP6P4b |
| Tibati | AfunF3_2 | 8546959 | 8553509 | Deletion | 6.5kb | 1/1 | Spanning CYP9a and b intergenic region |
| Tibati | AfunF3_2 | 8538070 | 8541606 | Deletion | 3.5kb | 0/1 | Spanning AFUN015973 and P450 AFUN008357 |
| Elende | AfunF3_2 | 8560655 | 8563261 | Duplication | 2.4kb | 0/1 | Spanning partial CYP6P4a and CYP6P4b |
| Elende | AfunF3_2 | 8536340 | 8539804 | Duplication | 3.5kb | 0/1 | Spanning partial 2X Carboxylesterases |
| Elende | AfunF3_X | 8358865 | 8373797 | Duplication | 14kb | 0/1 | Spanning CYP9K1 and AFUN007545 intergenic |
| Elende | AfunF3_2 | 8560530 | 8562886 | Deletion | 2.3kb | 0/1 | Spanning partial CYP6P4a and CYP6P4b |
| Elende | AfunF3_2 | 8546959 | 8553509 | Deletion | 6.5kb | 1/1 | Spanning CYP9a and b intergenic region |
| Elende | AfunF3_2 | 8560530 | 8562886 | Deletion | 2.3kb | 0/1 | Spanning partial CYP6P4a and CYP6P4b |
| Elende | AfunF3_X | 8347591 | 8377479 | Inversion | 29kb | 0/1 | Spanning CYP9K1 and AFUN007545 intergenic |
| Elende | AfunF3_X | 8356839 | 8405693 | Inversion | 48kb | 0/1 | Spanning entire AFUN007545 and next genes |
| Njombe | AfunF3_2 | 8536201 | 8539765 | Duplication | 3.5kb | 0/1 | Spanning partial 2X Carboxylesterases |
| Njombe | AfunF3_2 | 8527362 | 8534289 | Duplication | 6.9kb | 1/1 | Spanning entire CYP6AA1 and partial CYP6AA2 |
| Njombe | AfunF3_X | 8358940 | 8373624 | Duplication | 14kb | 1/1 | Spanning CYP9K1 and AFUN007545 intergenic |
| Njombe | AfunF3_2 | 8560518 | 8562882 | Deletion | 2.3kb | 0/1 | Spanning partial CYP6P4a and CYP6P4b |
| Njombe | AfunF3_2 | 8546959 | 8553509 | Deletion | 6.5kb | 1/1 | Spanning CYP9a and b intergenic region |
| Njombe | AfunF3_2 | 8561553 | 8564078 | Deletion | 2.5kb | 0/1 | Spanning partial CYP6P4a and CYP6P4b |

**Legend:** SV means structural variant; GT stands for genotype.

Two major deletions were identified across all populations: a 2.3 kb consistent deletion within the rp1 locus spanning partial *CYP6P4a* and *CYP6P4b* genes beside the 2.4 kb duplication found, and a 6.5 kb fixed deletion, an insertion in FUMOZ, observed in various localities across Africa [15]. The former was heterozygote with high quality across all populations. The latter (absence of 6.5kb insertion), although present in all four locations, exhibited poor heterozygote quality (Table 1, S5 Table, S6 and S7 Figs). Additionally, a unique 2.5 kb deletion with high heterozygote quality was exclusive to the Njombe population spanning the partial *CYP6P4a/b* paralogues (Table 1, S5 Table and S7 Fig).

In terms of inversions, multiple events of varying segment sizes, such as the 29 kb and 48 kb inversions downstream of the *CYP9K1* gene, were identified. These inversions were heterozygote in Gounougou and Elende populations but absent in other populations, including laboratory strains (Table 1 and S Table). Table 1 and supplementary tables provide detailed information on coverage depth, supporting reads, sequence quality, and heterozygote quality for each identified structural variant, aiding in a comprehensive understanding of their association to *An. funestus* adaptation to insecticide selection pressures or other ecological factors. The consistent patterns observed across populations underscore the importance of these structural variants in the evolutionary dynamics of mosquito adaptation to ecological pressures and/or responses to environmental pressures such as insecticide resistance.

Moreover, we conducted coverage analysis around both the rp1 and CYP9 loci using custom scripts. Coverage depth analysis involves computing the number of reads aligned to each genomic position, followed by normalization based on mean coverage depth for comparability. Summarization of normalized coverage within non-overlapping 1000-bp windows aids in identifying patterns of potential copy number variations across genomic regions and populations. The results revealed distinct patterns around these genomic regions. Around the rp1 locus, where the two carboxylesterases and 10 major cytochrome P450 genes are found (indicated between the two blue vertical lines in (S9 Fig), no evidence of apparent copy number variations was observed across all populations (S9 Fig). This finding suggests a stable copy number in this region, indicating the absence or the presence at low frequency of significant structural alterations that could impact the associated genes which is in line with heterozygote duplications found in this region.

Conversely, around the *CYP9K1* region where a TE was inserted upstream the gene, we identified evidence of copy number variations, exclusively in the Njombe population (S10a Fig, depicted between the two blue vertical lines). This observation implies the potential existence of at least one additional copy of the segment/gene on one of the two chromosomes, as the normal copy number is expected to be two (S10a Fig). Notably, other populations did not exhibit evidence of copy number variations within this specific region. However, there was apparent evidence of potential copy number variations downstream of *CYP9K1* gene, where several chromosomal inversions of substantial length were identified (S10 b-f Figs).

The absence of copy number variations around the rp1 locus in Elende, Tibati and Gounougou implies its relative stability across populations, contrasting with the dynamic nature of the *CYP9K1* region in Njombe population, warranting further investigation into the functional implications of these structural variations.

## Discussion

This study aims at deciphering patterns of genetic structure amongst populations of the major malaria vector *Anopheles funestus* across different eco-climatic landscape in Cameroon while identifying genomic signatures underlying adaptive responses to insecticides. It revealed a genetic subdivision associated with the eco-geographical patterns with northern populations exhibiting extensive gene flow and similar signatures of selective sweep associated with resistance different to southern populations.

### The population structure patterns in *An. funestus* populations across Cameroon provide essential insights is subdivided along eco-climatic lines

The population structure of malaria vectors is a critical factor in understanding the spread of insecticide resistance haplotypes and guiding effective implementation of intervention strategies like gene drive technology. The nation-wide analysis of *An. funestus* populations presented in this study provides valuable insights into their evolutionary history, population dynamics, and adaptation across diverse environmental contexts. The observed low *F_ST_* values and the PCA pattern between Gounougou and Tibati (Northern) signify extensive gene flow between these populations. The higher *F_ST_* values between Northern and Central (Elende) and Littoral/Coastal (Njombe) populations suggest weaker gene flow between these regions supported by the PCA pattern clearly clustering them separately. These findings align with studies conducted across Africa, which show restricted gene flow between *An. funestus* populations, both between different regions [15,32] and within regions of the same country [32,33]. Similar restricted gene flow has been reported in *An. gambiae* s.l. populations, notably in Southern Ghana and Burkina Faso [34,35], as well as in the Asian vector *Anopheles minimus* in Cambodia [36]. The Principal Component Analysis (PCA) further support these findings, revealing separate *An*. *funestus* population clusters in broad agreement with their respective eco-geographical regions emphasizing the role of geography in shaping genetic relationships. This study provides the foundation for future gene drive implementation in Cameroon, considering both biological, eco-geographical and ethical factors in *An. funestus* populations.

### Variability in evolutionary selection signals among *Anopheles funestus* populations in Cameroon is associated with insecticide resistance

The four *An. funestus* populations exhibit both common and varying signals of evolutionary selection dispersed throughout their genomes, particularly on chromosome 2R (CYP6/rp1) and X (CYP9). The rp1 locus houses two carboxylesterases and ten major cytochrome P450 genes, including the paralogues *CYP6P9a/b* and *CYP6P4a/b*, known for breaking down pyrethroids in mosquito populations from Southern and Western Africa, respectively [30,37–39]. Additionally, the *CYP9K1* gene within the CYP9 cluster, previously found to be highly overexpressed in Uganda *An. funestus* populations and involved in pyrethroids metabolism, contributes to the observed signals on X chromosome [17,40,41]. The differentiation in the rp1 signal observed in *An. funestus* populations in Elende, Tibati, and Gounougou (Central and Northern regions) contrasts with the stronger CYP9 signal in the Njombe population (Coastal region), indicating localized selection pressures. This contrasting pattern may arise from varying intensities and types of insecticide usage, leading to distinct selection forces across these populations. The stronger selection on the rp1 locus in areas like Tibati and Elende, could be attributed to specific insecticide usage more prevalent in the Northern and Central regions, coupled with distinct ecological or ecoclimatic factors. This pattern may signify the adaptive evolution of these populations in response to localized insecticide selection pressures. Additionally, beside to the intense selection force in Njombe, the limited gene flow between Njombe (Coastal) and Northern populations may contribute to the stronger CYP9 signal in Njombe. For the first time in Cameroon, beyond the rp1 locus, this study identifies another signal of selection located on the sex-linked X chromosome, with a stronger impact in the Njombe (Littoral/Coastal) compared to other populations. This suggests that resistance in Cameroon is not confined solely to the rp1 locus but extends to other genomic regions, highlighting the complexity and adaptability of resistance mechanisms. Comparable findings were observed in Uganda populations using targeted enrichment sequencing, revealing the emergence of both rp1 and CYP9 signals [40]. The identified signals indicate a robust adaptive and evolutionary response to environmental stimuli, particularly related to detoxification pathway. This adaptive response is potentially driven by sustained pesticide usage in the agricultural practices prevalent in these localities, which host major agricultural activities in Cameroon. Additionally, the extensive use of pyrethroid, organophosphate, and carbamate insecticides in agriculture likely contributes to the development of insecticide resistance in mosquito populations throughout these regions. Another signal of genetic differentiation was consistently observed in all six comparisons at the terminal end of the X chromosome, annotated with diacylglycerol kinase (DGK). This genomic region was detected to be under positive selection in *An. gambiae* and *An. coluzzii* in western Burkina Faso [35,42]. Research conducted in *Drosophila melanogaster* and *Caenorhabditis elegans* has indicated potential involvement of diacylglycerol kinase (DGK) in the adaptation to environmental changes [43,44]. The Drosophila retinal degeneration A gene (rdg A), which is an orthologue of the DGK gene, has been implicated in light signaling [44,45]. This probably suggests that if light signaling is regulated similarly in mosquito species, the DGK gene may play a role in the regulation of vision in *An. funestus* which could impact the mosquito behavior. However, future studies should aim to establish the direct or indirect contribution of this gene to the adaptive response to insecticides in these populations.

This study provides evidence of the evolving nature of *An. funestus* populations across Cameroon, likely driven by persistent and intense pressure from the use of agricultural insecticides. All four populations exhibit signs of selective sweeps and likely experienced recent population expansions with presence of rare alleles. This is supported by evidence of signatures of positive selection associated with reduced genetic diversity across all four populations, with stronger effects observed at the rp1 locus in Elende and Tibati populations compared to Njombe and Gounougou populations. Prior investigations reported emerging genetic differentiation, compelling evidence of selection, and reduced genetic diversity at the QTL rp1 locus in the *An. funestus* population from Malawi between 2002 and 2014 but not in 2014 in Mibellon locality which is in proximity with Tibati, in the Northern Cameroon [32]. The valley of reduced diversity found in our study represents a typical signature of a selective sweep, similar to what was previously described in the Southern African (Malawi and Mozambique) *An. funestus* populations [15,27]. This finding aligned to previous studies that identify positive selection around target site mutations (i.e. kdr) conferring resistance in many mosquito vectors species except *An. funestus* [46–50]. To comprehensively understand the dynamics of resistance-related loci and monitor their spread, expanded genomic surveillance is crucial across other regions in Cameroon. This approach will guide efforts aimed at malaria elimination in the country.

### Genetic variants including replacement polymorphisms are potentially driving insecticide resistance in *Anopheles funestus* across Cameroon

Selection pressure can lead to the increase in frequency of novel genetic variants, facilitating the adaptation and survival of malaria vectors. This is evident through the identification of non-synonymous single nucleotide polymorphisms (ns-SNPs) at varying frequencies across various populations in this study. Capitalizing on the strong evolutionary selection affecting metabolic resistance genes (both the P450-based rp1 and CYP9 loci), this study sought to explore specific genetic variants associated with the adaptive responses observed in these populations.

The *An. funestus* populations exhibit a multiallelic and multigenic pattern of ns-SNPs across the quantitative trait locus (QTL) rp1 and the CYP9 loci. The most significant novel ns-SNPs originate from *CYP6P2* (R346C), *CYP6P9a* (E91D and Y168H), *CYP6P9b* (V392F, V359I), *CYP6P5* (V14L) and *CYP6AA2* (V3A, S498L and P35S) genes spread around the rp1 locus, approaching fixation or already fixed in Elende and Tibati populations, displaying moderate to high frequencies in Gounougou, and occurring at lower frequencies in the Njombe population as well as in FANG fully susceptible strain. These genetic variations suggest a dynamic interplay between the genetic makeup of these populations and the selective pressures imposed by insecticide usage. These results align with the observed pattern of genetic differentiation, indicating a greater impact of rp1-based genetic variants in Northern and Central populations and lower impact in the Njombe Coastal population. Another fact is that the rp1 locus-based genetic variants exert a varying degree of impact on different *An. funestus* populations, suggesting potential regional differences in the selection pressure and adaptive responses to insecticide resistance. The multiallelic and multigenic pattern observed in these replacement polymorphisms suggests that they likely underwent collective selection over time and therefore, some of these SNPs may share the same haplotype. Unfortunately, the PoolSeq approach utilized in this study had limitations, preventing the conduct of comprehensive analyses such as haplotype network, clustering, and linkage disequilibrium assessments. This complexity poses a challenge in pinpointing specific haplotype driving insecticide resistance to specific regions. The presence of a 4.3 kb transposable element (TE) insertion, identified within the rp1, precisely in the intergenic region between *CYP6P9b* (downstream) and *CYP6P5* (upstream), contributing to pyrethroid resistance adds another layer of complexity, suggesting that it may have an additive impact combined with those ns-SNPs within the rp1. Other novel ns-SNPs found within the rp1 at moderate to high frequencies included the known SNPs presented above on *CYP6P4a/b* paralogues and the SNPs on *CYP6P1* (I17T), P450 *AFUN019478* (F60S and F371Y). Except for H169R, all mutations on *CYP6P4a* and *b* were recently reported at very low frequencies (<15%) in *An. funestus* populations in the Democratic Republic of the Congo using amplicon sequencing, potentially indicating a widespread geographical distribution of resistance-associated variants [51]. The *GSTe2* (L119F) known point mutation conferring DDT and permethrin resistance in Cameroon (56) and in Benin (57) was identified at moderate to high frequencies across locations suggesting contribution of this mechanism in resistance with varying impact across regions. Interestingly, a new *GSTe2* variant (F120L) was identified next the previous one at moderate frequencies potentially meaning that they co-occurred together, a hypothesis that need to be validated in future studies. Additionally, It was also different from the novel mutation, K146T, identified in the Democratic Republic of the Congo [51]. These findings highlight the diverse repertoire of *GSTe2* mutations in *An. funestus* populations, suggesting ongoing evolution and adaptation to insecticide exposure. The presence of novel mutations further underscores the complexity of insecticide resistance mechanisms, necessitating continuous surveillance to inform targeted control strategies. A search for SNPs on *An. funestus* Voltage Gated Sodium Channel (VGSC) revealed no knockdown mutations for this species in Cameroon, unlike *An. gambiae*, where the first genetic marker of resistance against pyrethroids, kdr-L1014F (or L995F), was identified almost 25 years ago [52]. The two identified SNPs, D1156Y and L1988I, were found at low frequencies in all populations (<20%). The apparent lack of a significant role for kdr in adaptive responses to selection pressure underscores the predominant contribution of other mechanisms such as metabolic mechanism in *An. funestus* populations.

In contrast, the most significant and only CYP9-based ns-SNP results in a glycine-to-alanine substitution at position 454 on the *CYP9K1* gene. This variant has been reported near fixation in the Njombe population and remains under selection in the Elende, Tibati, and Gounougou populations [41], aligning with the pattern of high differentiation observed in Njombe compared to elsewhere. This suggests that the *CYP9K1* gene could be a primary genetic factor associated with the adaptive response in the Littoral/Coastal *An. funestus* population. In Southern Ghana, a recent study on *An. gambiae* sibling species reported the presence of a ns-SNP different from the current point mutation. Specifically, the N224I mutation was identified, indicating that it is undergoing intense selection and is present at a high frequency (∼60%) in the *An. gambiae* sl population in that region [34]. Our finding suggests that different genetic variations are under selective pressure in distinct geographic locations in diverse species, highlighting the dynamic nature of mosquito populations and their responses to environmental factors.

### Candidate signatures of complex genomic evolution potentially driving insecticide resistance in *Anopheles funestus* across Cameroon

We have documented a series of complex genomic anomalies within the CYP6 rp1 cluster, likely linked to the previously reported selective sweeps driven by selection pressure in *An. funestus* populations across Cameroon. These complex features encompass a novel duplication of a 2.4 kb segment containing partial paralogous *CYP6P4a/b* sequences and a 3.5 kb fragment partially spanning the two rp1-associated carboxylesterases. Several studies have investigated the overactivity of metabolic resistance genes, such as *CYP6AA1* with supporting evidence of duplications occurring in *An. funestus* populations across Africa [53]. The findings align with previous evidence of *CYP6AA1* gene duplications observed in *An. coluzzii* [54,55] and *An. gambiae* populations from East and Central Africa resistant to deltamethrin. In the latter populations, the copy number variation (CNV) *Cyp6aap_Dup1* has been identified as being linked to deltamethrin resistance and has demonstrated rapid dissemination in these regions [56]. These results deviate from the anticipated copy number pattern, as there is no evidence of an increased copy number within the rp1 locus. This inconsistency may be explained by the presence of deletions within these duplications, some of which are larger in size, spanning entire genomic segments or by the genotype pattern who indicates that these duplications are heterozygote and therefore still under selection in these localities. Common deletions for all four populations included a 2.3kb of partial *CYP6P4a*/*b* sequences and a 6.5kb intergenic deletion between *CYP6P9a*/*b*, an insertion in the FUMOZ reference genome previously reported in Malawi and across other African countries [15,17]. While the Southern Africa insertion was absent in all four populations, other duplications were identified as unique to the Njombe and Elende populations, confirming the contrasting pattern of genomic evolution in the Central and Littoral (Coastal) regions compared to the Northern region. Specifically, a 14 kb duplication was found to be already fixed in Njombe but not in Elende, where it is still under selection in line with pattern of copy number variant. This duplication spans the intergenic region of *CYP9K1* and the next gene, *AFUN007545*, aligning with the pattern of increased copy number previously presented around this region in Njombe.

We detected evidence of a selective sweep at the CYP9 locus, encompassing the *CYP9K1* gene in the Njombe population, characterized by an increased copy number and high coverage depth flanking this genomic locus. This was unique to the Njombe population, not observed in other locations. Interestingly, this population exhibited at least one additional copy of the gene, aligning with the observed genetic differentiation pattern. Additionally, within the coding region of the *CYP9K1* gene, we identified a fixed ns-SNP (G454A). This specific SNP, previously fixed in the Ugandan population [40], has been identified as a major driver of pyrethroid resistance evolution in Mibellon, Cameroon with overexpression of the corresponding gene and fixation in Njombe natural population while still under selection elsewhere [15,41]. Furthermore, upstream of the *CYP9K1* gene, a transposon insertion similar to the one detected in Uganda [15] with incomplete assembled sequence due to the long length, appeared to be consistently present in the Njombe population compared to other localities. This observation suggests possible evidence of gene flow between Eastern and Central Africa. The overexpression of this gene was associated with the presence of an identical transposable element (TE) and gene duplication, causing an extensive selective sweep in *An. funestus* population in 2014 in Uganda [17,57], which we hypothesized could be the same scenario happening in Njombe Coastal *An. funestus* population. Future studies should also prioritize unravelling the complete sequence of this substantial transposable element (TE) employing third-generation sequencing techniques.

## Conclusion

This study offers conclusive evidence of distinct patterns of selective sweeps acting on two major loci, rp1 and CYP9, accompanied by genetic variants and complex genomic alterations in *An. funestus* populations across Cameroon. These adaptations have emerged in response to varying selection pressure in *An. funestus* populations across four agricultural settings in Cameroon. Future investigations should expand genomic surveillance across the Central African regions to track the dynamics of resistance-related loci, haplotypes and guide efforts toward malaria elimination, considering the complex and multigenic nature of insecticide resistance in *An. funestus* populations. Further investigations are warranted to elucidate the functional consequences of these novel replacement polymorphisms and these genomic complex features, their impact on resistance phenotypes, and their potential role in shaping the evolutionary dynamics of insecticide resistance in malaria vectors.

## Models

### Ethical considerations

This study underwent review and approval by the National Ethics Committee for Health Research (CNERSH) of Cameroon, with the identification number 2021/07/1372/CE/CNERSH/SP.

### Sample collection

*An. funestus* mosquito samples were collected from four distinct eco-geographical locations across Cameroon, including Gounougou (9°03’00’’N, 13°43’59’’E) in the north and Tibati (6°27‘57”N; 12°37‘30”E) in Central-North, Elende (3°41’57.27’’N, 11°33’28.46’’E) in the central, and Njombe (4°35‘00”N, 9°40‘00”E) in the littoral/coastal regions (Fig 9). These regions are characterised by two main climates: the equatorial and sub-tropical climate found in the Littoral/Coastal region (Njombe) and Centre region (Elende), and the tropical climate present in the Northern regions (Gounougou and Tibati).

**Figure 9:**
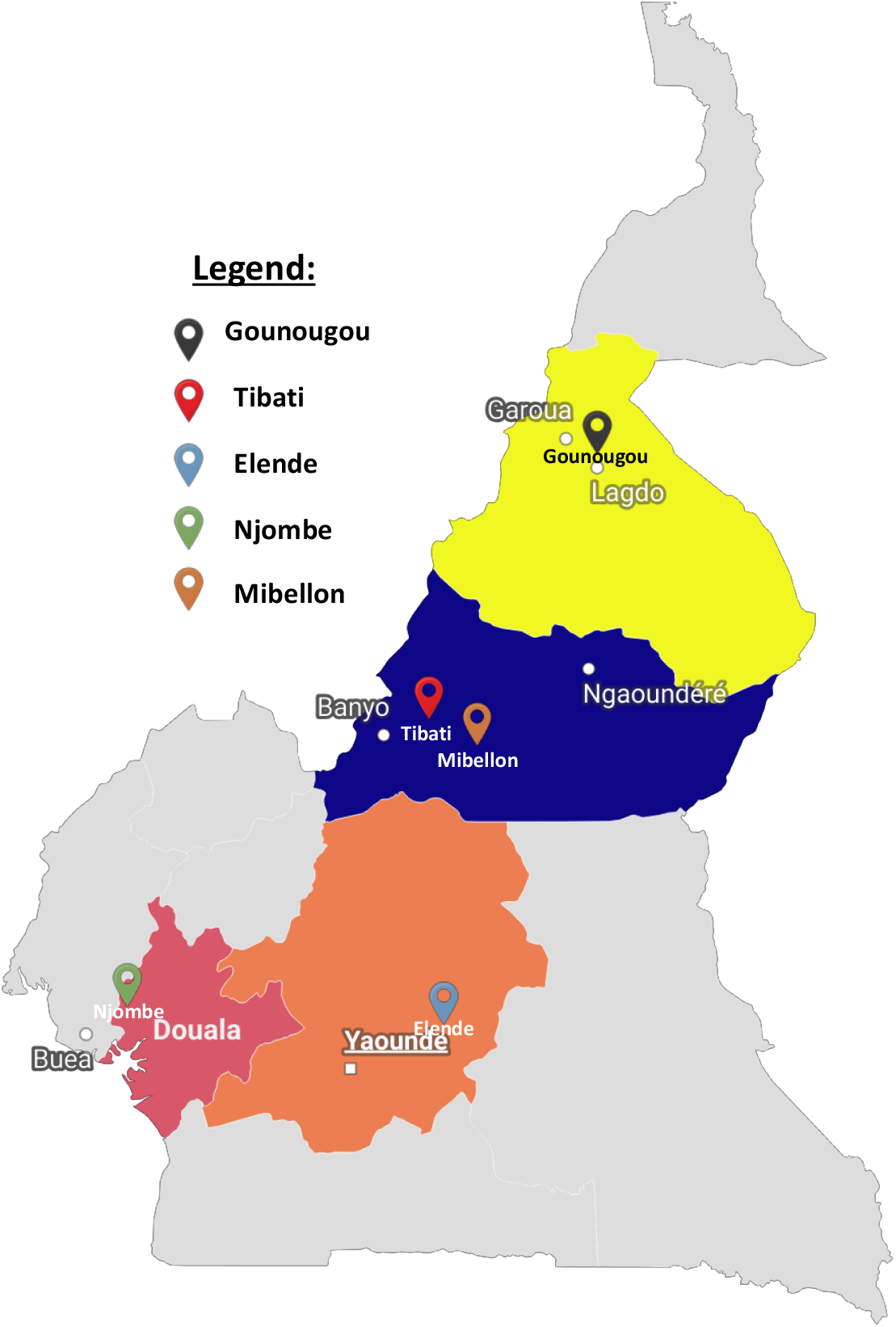
Study sites map.

Gounougou, located near the Benoué River, houses a significant hydroelectric dam providing electricity and supporting irrigation for 15,000 hectares of downstream crops. The presence of vegetation along the riverbanks and a large rice field creates favourable conditions for the primary malaria vector, *Anopheles funestus*, with extensive insecticide use, including pyrethroids and DDT [58]. Tibati is in the Adamawa region of Cameroon located between the north and south ecoclimatic zones in Cameroon. Elende is a rural village located about 2 km from the Yaoundé-Nsimalen International Airport. Elende relies on subsistence farming for crops like cassava, cocoa, and vegetables, significantly enhanced by pesticide application [59]. Njombe-Penja is located in the Moungo department in the Littoral region. The town is renowned for its expansive plantations, The cultivation spans various crops, including cocoa, pepper, and exotic flowers, covering a combined production area of 5,000 hectares. Intensive pesticide use may contribute to the selection of resistant phenotypes in malaria vectors. *An. gambiae* and *An. funestus* are the principal malaria vectors responsible for a substantial malaria transmission in these eco-geographical regions [58,59]. *An. funestus* populations from these ecoregions have been reported to exhibit high resistance intensity to both types I and II pyrethroid insecticides. In all localities, collection with consent of blood-fed adult female *Anopheles funestus* mosquitoes resting indoors occurred between 06:00 a.m. and 10:00 a.m., utilizing torches and Prokopack electric aspirators (John W Hock Co, Gainesville, FL, USA). Additionally, the genomic study included two laboratory reference strains and one field strain collected at Mibellon (North-West) in 2014: FUMOZ from Southern Mozambique, exhibiting multiple insecticide resistance, and FANG from Southern Angola, a fully susceptible strain used for validating positive selection signals [60]. This Mibellon 2014 sample served as a geographically closer negative control than the FANG sample, because there was no evidence of selection detected in this population [15].

### DNA extraction, sequence library preparation and pool whole genome sequencing

PoolSeq GWAS experiment used one biological replicate consisting of 40 individuals per pool in each locality. Genomic DNA was extracted from individual female mosquito using the DNeasy Blood and Tissue kit from Qiagen [61]. Molecular species identification was subsequently performed as previously described by [62]. The DNA samples were purified using Rnase A and quantified using a Qubit instrument then pooled in pool of 40 individuals per replicate in equal amounts of DNA. Library preparation, quality control and pair-end (2 × 150 bp) whole-genome sequencing with NovaSeq Illumina sequencer was carried out to a target 50x coverage by Novogene (Cambridge, United Kingdom).

### Computational analyses

#### Quality control of fastq files, mapping, filtering and creation of synchronized format

The pipeline for the analyses of our PoolSeq dataset is available on GitHub repository via https://github.com/Gadji-M/PoolSeq_OMIcsTouch. Briefly, quality control procedures were executed using FastQC (fastqc, 2019), and the results were subsequently piped into Multiqc for aggregation and visualization of the final outcomes [63]. Alignment was conducted using the reference genome of *An. funestus* (VectorBase-61_AfunestusFUMOZ_Genome.fasta) sourced from VectorBase (https://vectorbase.org/vectorbase/app/downloads/release-61/AfunestusFUMOZ/fasta/data/). in ’bwa mem’ [64], output was then piped using "samtools view -Sb -" to generate a BAM file containing only reads with a quality Phred score greater than 10 (-q 10). The alignment files in BAM format underwent sorting, marking, and deduplication processes using the PICARD tool (https://broadinstitute.github.io/picard/). The synchronized file function is the principal input files for PoPoolation2 [65], and therefore is key for the downstream analyses. It contains the allele frequencies for each population (pool) at every base of the reference genome in a condensed format. To generate a synchronized file, an initial multiple pileup file (mpileup) was created by amalgamating data from all population pools. The "Samtools mpileup -B -Q 0" command was employed to produce a unified mpileup file. Subsequently, the generated mpileup file underwent processing using the "java -ea -jar mpileup2sync.jar --threads 40" command in PoPoolation2 to yield the synchronized output file [65].

#### Tracing selective sweeps with population-based genomics analysis

The synchronized file was used in downstream analyses for the estimation of various population genomic estimators. Indeed, a bespoke popoolation2-based shell script, "Fst_sliding_windows.sh" [65] was employed to assess precise allele frequencies and pairwise *F_ST_* genetic differentiation across various population pools. The script "perl snp-frequency.pl" with synchronized file as input, facilitated the computation of exact allele frequency differences, with parameters set at --min-count 2, --min-coverage 10, and --max-coverage 5%. Principal Component Analysis (PCA) for population structure analysis utilized output from "perl snp-frequency.pl". Additionally, pairwise average *F_ST_* for all possible comparison among all pools was determined using the custom script "Fst_sliding_windows.sh" across sliding windows ranging from 5kb to 100kb. The formatted *F_ST_* files and relevant comparisons were visualized using the ggplot2 package in R [66]. Additionally, Genome-wide *F_ST_* and heterozygosity between populations were computed using poolfstat [67].

To showcase evidence of selective sweeps among populations/phenotypes, Genome-wide Tajima’s D values and theta pi nucleotide diversity (π) were computed for each pool sample in overlapping windows of 50kb using grenedalf (https://github.com/lczech/grenedalf). All the genomic regions detected with high signal of genetic differentiation and potentially under strong evolutionary processes were analysed in detail by performing fine-scale resolution analysis zooming into them in overlapping windows of 1 kb moving in steps of 0.5 kb via grenedalf (https://github.com/lczech/grenedalf).

#### Variant calling and filtering

A comprehensive variant calling was done with VarScan command line-based tools. It employs heuristic and statistic thresholds based on user-defined criteria to call variants using Samtools mpileup data as input to detect SNPs/indels in individual and pooled samples [68]. Outputs SNPs obtained post VarScan were filtered to just retain bi-allelic variants. The final SNPs and indels variants calling format (vcf) files were annotated and variants were predicted using SnpEff [69] which annotates and predicts the effects of genetic variants (such as amino acid changes). For that, *An. funestus* database was built and SnpEff was run with java -Xmx8g -jar snpEff.jar command line then the output file was filtered with SnpSift and bcftools [69,70].

#### Exploring complex genomic evolutionary signatures

Combination of computational tools including INSurVeyor [71] and Smoove (https://github.com/brentp/smoove) coupled to Integrative Genomic Viewer (IGV) were used to detect and visualized complex signatures of genomic anomalies including large duplications, large chromosomal inversions, large translocations and large indels (large SVs). Large structural variants were defined as variation of the DNA segment or gene in the genome including more than 1,000 bp. Large SVs analysis used the BAM alignment file for each *An. funestus* population. INSurVeyor used python programming language through "python insurveyor.py" command to detect all genomic alterations associated to insertions (large insertions) whereas smoove through conda, was used to detect additional structural variations (SVs) not detected by INSurVeyor such as inversions, deletions and duplications via "smoove call" command line. The identified variants were annotated using “smoove annotate” and filtered with bcftools filter [70].

The structural variants detected through computational approach were further visualized in IGV. The examination and visualization of BAM alignment files aimed to deduce genomic anomalies through consideration of four key metrics. Firstly, an increased coverage depth spanning specific gene clusters or genomic regions, indicating potential duplication with the presence of more than one copy of the genomic fragment or the insertion of a large DNA fragment. Secondly, the identification of read pairs with incorrect insert sizes within the genomic regions. Thirdly, the detection of abnormal relative read pair orientations suggesting a variety of structural variations, including duplication, intra or inter chromosomal translocation, indels, and chromosomal inversions. Lastly, the observation of multiple seemingly chimeric and discordant reads spanning putative breakpoints, either clipped or not, in the alignment, further contributed to the inference of genomic alterations.

## Supporting information

**S1 Fig. Pairwise *F_ST_* genetic differentiation between *Anopheles funestus* population from four eco-geographical settings and FANG highly susceptible laboratory strain.** A, B, C, D, E, F represent genome-wide comparisons between control versus FANG, Tibati versus FANG, Gounougou versus FANG, Elende versus FANG, Djombe versus FANG and FANG versus FUMOZ highly resistant populations, respectively.

**S2 Fig. Evidence of a 4.3kb transposon insertion between CYP6P9b and CYP6P5 is observed in *Anopheles funestus* populations across Cameroon.** The screenshot from Integrative Genomics Viewer (IGV) displays coverage depth and aligned reads for pooled template whole genome sequences of FANG, FUMOZ, Gounougou, Tibati, Elende, and Njombe. The red markers in the circle indicate the TE insertion, displayed with a characteristic pattern in blue vertical rectangle. The coverage depth plots reveal increased coverage downstream of the TE insertion in all populations. The grey rectangles separated by thin lines represent normal reads aligned in pairs, while green rectangles represent discordant reads.

**S3 Fig. Evidence of a substantial transposon insertion, with an unknown size, observed upstream of the CYP9K1 gene in *Anopheles funestus* populations across Cameroon.** The red markers in circle indicate TE insertion in all populations, characterized by a pattern as blue vertical rectangle, apparently more prevalent in Njombe population compared to others where it is still under selection. The coverage depth plots reveal increased coverage downstream of the TE insertion in the Njombe population but not in others, including FANG and FUMOZ. The grey rectangles represent normal reads aligned in pairs, while thick black rectangles represent discordant reads.

**S4 Fig. Evidence of a 2.4kb duplication, spanning partial CYP6P4a and b paralogues, is observed with an unusual insert size, represented by green rectangles separated with thin lines and framed in the blue boxe.** These duplications are present in Gounougou, Tibati, and Elende but absent in Njombe, FANG, and FUMOZ. A red framed box with reads of unusual sizes, separated by red thin lines, corresponds to a 2.3kb deletion found in all populations, while the orange framed box represents a 2.5kb deletion only found in the Njombe population. The drop in coverage is noticeable in the Njombe population coverage track. Grey rectangles, separated by thin lines, represent normal aligned reads. Overlapping duplications and deletions point to a complex genomic alteration that need further investigations.

**S5 Fig. Evidence of a 3.5kb duplication, spanning partial 2x carboxylesterases (rp1-based carboxylesterases), is represented by an unusual insert size as green rectangles, separated by a thin line and framed in a blue box.** This duplication is consistent in all field populations but with lower supporting reads in Njombe. It is absent in FUMOZ and has just two supporting reads in FANG. The grey rectangles present on the alignment track of FUMOZ represent normal aligned reads viewed in pairs and separated by a thin line.

**S6 Fig. Evidence of a 2.3 kb deletion, spanning partial CYP6P4a and b paralogues, is represented by an unusual large insert size in red rectangles, separated by a thin line and boxed with red boxes.** This deletion is more consistent in Gounougou, Tibati, and Elende but with lower supporting reads in Njombe. Conversely, another deletion of 2.5kb was only found in Njombe, overlapping with the 2.3kb deletion, shown in an orange box with a drop in coverage depth in this population. All these deletions were absent in FANG and FUMOZ. The grey rectangles present on the alignment track of FANG and FUMOZ represent normal aligned reads viewed in pairs and separated by a thin line.

**S7 Fig. Evidence of a 6.5 kb deletion (absence of insertion) between CYP6P9a and b (intergenic) paralogues is represented by an unusual large insert size in red rectangle, separated by a thin line and framed in red boxes.** This corresponds to an insertion in FUMOZ, which is absent in all our field Cameroon populations, including FANG. The grey rectangles represent normal aligned reads viewed in pairs and separated by a thin line.

**S8 Fig. Minor and Major allele frequencies spanning the entire rp1 and CYP9K1 loci in *Anopheles funestus* populations across Cameroon.**

**S9 Fig. Coverage analyses showing regions potentially affected by CNVs around the QTL rp1 locus in *An. funestus* populations across Cameroon.**

**S10 Fig. Coverage analyses showing regions potentially affected by CNVs around the CYP9 cluster in *An. funestus* populations across Cameroon.** Each dot represents a coverage window of 1000 bp.

**S1 Table. Alignment statistics of *An. funestus* PoolSeq whole genome sequencing data across Cameroon.**

**S2 Table. Coverage statistics of *An. funestus* PoolSeq whole genome sequencing data across Cameroon.**

**S3 Table. Missense and putative variants associated to major selective sweeps in *An. funestus* population across Cameroon.**

**S4 Table. Structural variants-based insertions associated to major selective sweeps in *An. funestus* populations across Cameroon.**

**S5 Table. Structural variants-based duplications and deletions associated to major selective sweeps in *An. funestus* across Cameroon.**

## Author contributions

CSW conceived and design the study. MG implemented the study design, executed the experimental work and analysed data. MG, JAKO, JH and CSW collaborated on data analysis, visualization, and biological interpretation of findings. MG, MT, MJW and LM provided laboratory resources, conducted field and experimental work. MG drafted the manuscript with assistance from CSW and contributions from all co-authors. CSW and BO provided supervision. All authors participated in reviewing and approving the final version of the manuscript for submission.

## Code availability

All the Codes used to analyse the data are available in the GitHub repository https://github.com/Gadji-M/PoolSeq_OMIcsTouch.

## Data availability

The datasets from the PoolSeq whole genome sequencing are accessible on the European Nucleotide Archive under accessions PRJEB76574.

